# Convergent Mitochondrial and Oxidative Stress Signatures in Spaceflight and Tauopathy: A Comparative Multi-Omics Reanalysis in Drosophila melanogaster

**DOI:** 10.64898/2026.10.16.759726

**Authors:** Marian Navarro, Jelard Aquino, King Shi, Henry Lin, Vahid Monfared, Joven Christopher Garcia, Allen Seylani, Maria Komędera, Julia Moutinho, Jessica Romils, Andrea Venosa, Anish Kelam, Vandreza Cardoso Bortolotto, Jingchun Chen, Matthew Gebre, M Windy McNerney

**Affiliations:** Florida International University, 11200 SW 8th Street, Miami, FL 33199, United States; Howard University, 520 W St NW, Washington, DC 20059, United States; Nevada Institute of Personalized Medicine, University of Nevada, 4505 S Maryland Pkwy, Las Vegas, NV 89154, United States; Johns Hopkins University, 3400 N. Charles Street, Baltimore, MD 21218, United States; Independent Researcher, Fremont, CA 94538, United States; Skeletal Biology Laboratory, School of Nutrition and Public Health, Oregon State University, Corvallis, OR 97331, United States; University of Central Florida, 4000 Central Florida Blvd, Orlando, FL 32816, United States; Cleveland Clinic, Ohio, USA, Internal Medicine Department; SWPS University, al. Adama Mickiewicza 3, 31-120 Kraków, Poland; Universidade Federal de São Paulo (UNIFESP), Rua Botucatu, 740, Vila Clementino, São Paulo, SP 04023-900, Brazil; School of Clinical Medicine, University of Cambridge, Cambridge Biomedical Campus, CB2 0SP, United Kingdom; Independent Researcher, Holbæk, Denmark, 4300; University of Virginia, 1827 University Avenue, Charlottesville, VA 22903; Universidade Federal do Pampa (UNIPAMPA), Rua Luiz Joaquim de Sá Britto, s/n, Promorar, Itaqui - RS, Brazil; Laboratory of Pharmacological and Toxicological Evaluations Applied to Bioactive Molecules (Laftambio Pampa) - Nutrition Department; Independent Researcher, Cypress, CA 90630, United States; MIRECC, VA Palo Alto Health Care System, 3801 Miranda Ave, Palo Alto, CA 94304, United States; Department of Psychiatry and Behavioral Sciences, Stanford University School of Medicine, 401 Quarry Rd, Stanford, CA 94305, United States

**Keywords:** Alzheimer’s Disease, Neurodegeneration, Oxidative Phosphorylation, Microgravity, Radiation

## Abstract

Mitochondrial dysfunction is a shared biological thread between terrestrial Alzheimer’s disease (AD) and in the unique stress environment of spaceflight, making mitochondrial pathophysiology a compelling target for comparative investigation. In AD, mitochondrial impairment is closely linked to defective oxidative phosphorylation, reactive oxygen species accumulation, calcium dysregulation, and neuronal injury, while spaceflight stressors such as microgravity and radiation disrupt mitochondrial homeostasis and increase oxidative stress. Understanding how the extreme stressors of spacelight affect mitochondrial function may provide insights into conserved mitochondrial stress responses and their implications for neurodegenerative diseases, including AD. Our study compares datasets from ground control Drosophila melanogaster RNA-seq with isolated radiation and microgravity exposure to discern conserved mitochondrial pathways and molecular signatures that may provide insights into AD mechanisms.

**Highlights:**

1. Progressive mitochondrial upregulation observed from Earth to microgravity
2. Tauopathy showed opposing downregulation of mitochondrial pathways
3. Shared ETC and ATP synthesis pathways connect microgravity and Tauopathy
4. Radiation induced minimal transcriptomic changes in spaceflight datasets
5. Mitochondrial upregulation may reflect compensation in response to microgravity

## 1. Introduction

Alzheimer’s disease (AD) currently affects over 7 million Americans and 33 to 38 million persons worldwide; projections estimate that this number will rise to 13.86 million individuals by 2060 [1, 2]. As the duration and frequency of human spaceflight missions increase, further research on how spaceflight stressors disrupt cellular homeostasis and mitochondrial function is needed to better understand and mitigate potential risks to crew health. Exposure to spaceflight stressors such as microgravity and cosmic radiation has been shown to cause immune system dysregulation, musculoskeletal atrophy, circadian rhythm disruption, and increased cancer risk [3-8]. Beyond these organ-system effects, comprehensive multi-omic analyses across tissues and organisms have identified mitochondrial stress as a central biological hub of the spaceflight response [9, 10]. This convergence is of particular interest given the well-established role of mitochondrial dysfunction in AD pathology and in age-related neurodegeneration more broadly [10-13].

While much research over the past 30 years has focused on removing β-amyloid (Aβ) plaques and tau neurofibrillary tangles (NFTs) to treat AD, it has become apparent that these targets alone may not fully explain the etiology or progression of the disease [14, 15]. Research has since expanded beyond plaques and NFTs to examine more upstream processes, leading to the development of the mitochondrial cascade hypothesis (MCH) [15-17]. Comparisons of transcriptomic data between individuals with and without AD have revealed differences in mitochondrial bioenergetics, particularly in the oxidative phosphorylation (OXPHOS) pathway and electron transport chain (ETC) [15, 16]. Additionally, experimental complex I deficiency has proven sufficient to induce AD-like molecular signatures that are reversible by targeted therapy, providing direct evidence that bioenergetic failure can contribute to processes upstream of pathology rather than solely representing a downstream consequence [18]. These disruptions lead to a buildup of reactive oxygen species (ROS) that contributes to Aβ accumulation, further impairing bioenergetic pathways and amplifying ROS production [15, 16, 19]. Due to the relatively low intrinsic antioxidant capacity of neurons,, ROS accumulation is thought to be a contributing factor to neuronal death in AD [20]. Additionally endoplasmic reticulum (ER) impairment in AD elevates cytosolic Ca²⁺, drives mitochondrial Ca²⁺ overload, and ultimately promotes cytochrome c release [21-23].

Growing evidence links specific spaceflight stressors to these same mechanisms. Simulated radiation exposure increases mitochondrial ROS production in rodents [24, 25], and these findings have been associated with cognitive decline and proteomic changes previously implicated in early AD pathology, including markers such as FAM162a and CST3 [8, 26, 27]. Simulated microgravity produces comparable impairments, with rats showing elevated mitochondrial ROS generation [28]. Disruption of Ca²⁺ homeostasis is a feature of AD pathology and has likewise been observed following exposure to spaceflight-relevant stressors. Studies have linked altered expression of ER and mitochondrial proteins, including STIM2 and MICU3, to increased ROS production, impaired ATP synthesis, reduced mitochondrial membrane potential, and mitochondrial Ca²⁺ overload leading to neuronal apoptosis [29-31]. This dysfunction is not confined to neurons, as astrocytes and microglia shape AD progression through inflammatory signaling and metabolic coupling. Astrocytic mitochondrial dysfunction, impaired mitophagy, and disrupted lipid metabolism can promote neuroinflammation and neurodegeneration, while APOE4 alters mitochondrial dynamics and OXPHOS [32-37]. Although astrocytes can transfer functional mitochondria to stressed neurons and restore bioenergetic function, this capacity may be limited by mitochondrial damage and disease progression [17, 38-44]. Radiation may further compromise glial support through ROS overproduction, Ca²⁺ dysregulation, altered connexin 43 signaling, impaired mitophagy, mitochondrial DNA release, and senescence, ultimately weakening metabolic support for neurons and oligodendrocytes [45, 46]. Together, these findings suggest that spaceflight stressors may disrupt neuronal and glial mitochondrial processes relevant to AD and the mechanisms described by the MCH.

Most studies examine mitochondrial dysfunction, AD pathology, and spaceflight effects either in isolation or in pairwise comparisons. Spaceflight effects are rarely delineated into their separate constituent factors such as radiation and microgravity, and glial contributions to mitochondrial stress are typically addressed apart from neuronal ones despite their metabolic interdependence. *D. melanogaster* offers a tractable system for addressing these gaps. Specifically, the findings that core mitochondrial bioenergetic machinery is highly conserved, astrocyte-like glia perform conserved metabolic coupling with neurons, and ground-based and flight datasets permit radiation and microgravity to be examined as separable exposures within a single organism. Our study uses *D. melanogaster* RNA-seq and proteomics datasets from three independent studies to compare the effects of isolated spaceflight stressors on mitochondrial pathway enrichment across radiation and microgravity conditions relative to ground controls, and to relate these findings to conserved neuronal and glial mitochondrial mechanisms implicated in AD pathology. By identifying conserved mitochondrial responses to spaceflight-relevant stressors, this work aims to explore how environmental stress influences pathways associated with neurodegeneration, as well as the potential relevance of spaceflight biology to AD.

## 2. Methods

### 2.1 Data Acquisition and Sample Information

Datasets were required to meet the following criteria for inclusion in the analysis: (1) *D. melanogaster* as the primary model organism, (2) whole brain as the assayed tissue, and (3) bulk RNA sequencing data available. Datasets were then categorized as either belonging to the spaceflight condition, the AD condition, or the radiation exposure condition [Figure 1].

**Figure 1.**
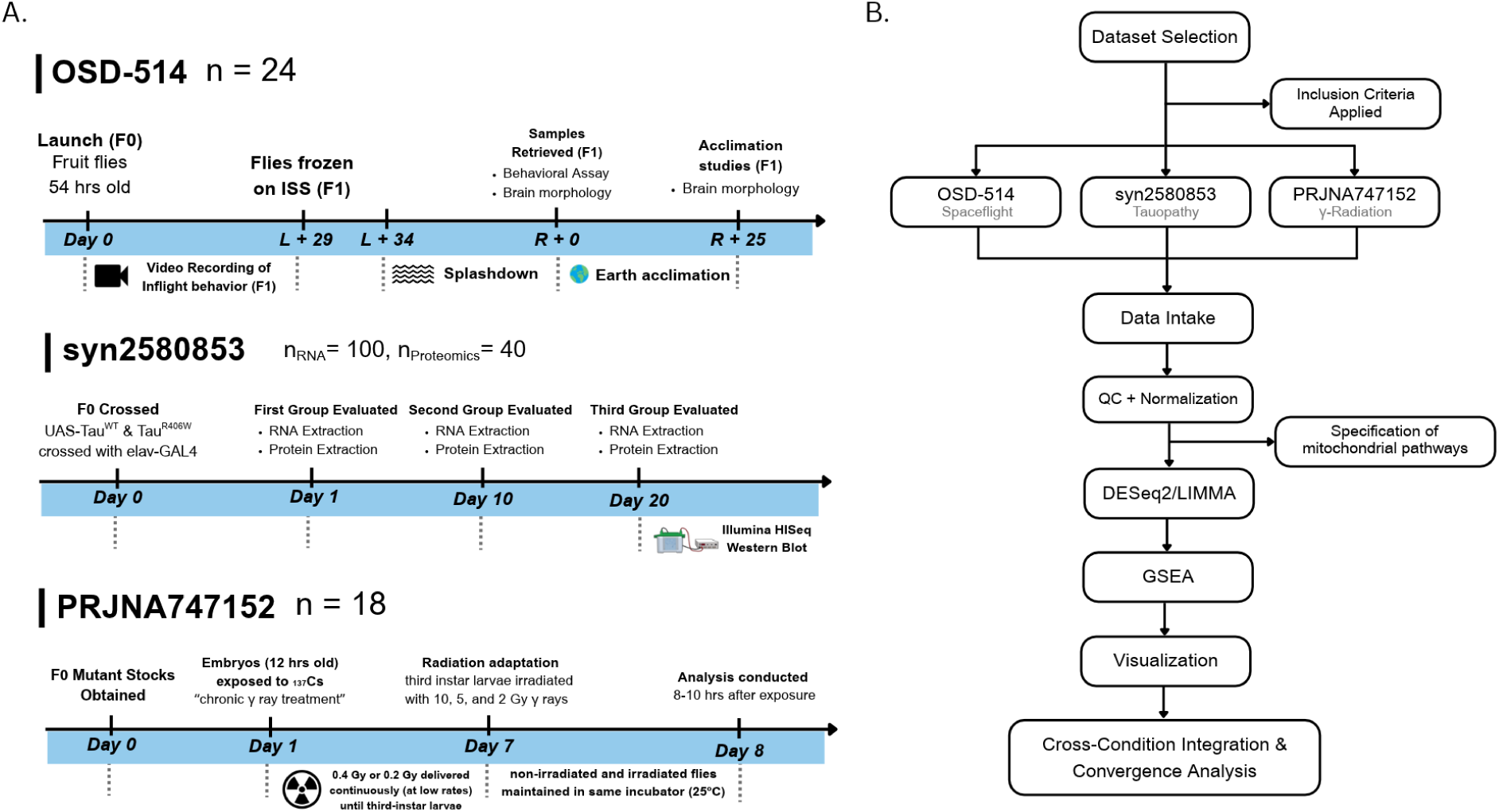
(A) A visual representation of the three selected datasets and the experimental conditions they underwent (B) The unified processing pipeline that was applied to all three datasets, from dataset selection to comparative figure generation.

The samples in OSD-514 were fruit flies aboard the ISS on the MVP-Fly-01 mission. The flies were divided into three conditions: Earth Control (authentic 1g), spaceflight with microgravity, and spaceflight with simulated 1g (centrifuged). Each group contained an F1 generation of four males and four females with wild-type genotypes. Samples were 6-12 days old at the time of sacrifice and tissue collection [47].

The samples in syn2580853 were fruit flies divided into three main genotype groups: Control (no Tau expression), wild-type human Tau expression (TauWT), and mutant Tau expression (TauR406W). The TauWT group expressed non-mutant human Tau and modeled early stage Tauopathy, while the TauR406W group expressed a disease-associated mutant that has been linked to neurodegeneration. Each group was sampled at several time points (1 day, 10 days, and 20 days) to observe progression over time [48].

The samples in PRJNA747152 were fruit flies exposed to ionizing radiation to study neural responses to radiation-induced stress. The flies were divided into two conditions: Control flies with no radiation exposure and radiation-exposed flies. All flies were adult wild-type males, and brain tissue was dissected and pooled for each RNA-seq sample [49].

### 2.2 Preprocessing and Quality Control

#### 2.2.1 RNA-seq

The three RNA datasets were preprocessed in R version 4.5.1 (2025-06-13) using a common pipeline, with dataset-specific deviations outlined below. Per-sample gene-level count files were compiled into a single count matrix, with missing genes assigned a count of zero wherever applicable, and sample metadata was parsed from filenames or accompanying metadata tables. Library sizes were computed per sample and inspected to identify potentially under-sequenced libraries. Genes were included in the analysis if they had ≥10 counts in ≥20% of samples. Filtered counts were used to construct a DESeq2 [50] dataset, and counts were variance-stabilized (VST, blind = TRUE) for exploratory analysis. Principal component analysis (PCA) was then performed on the VST-transformed data to assess sample clustering by condition, genotype, timepoint, and/or sex.

Two dataset-specific deviations from this pipeline should be noted. First, because OSD-514 RSEM outputs “expected counts” as fractional values, raw genes results files were imported with tximport (v1.36.1) rather than as a simple count matrix. This approach generates length-aware normalization offsets used directly by DESeq2 and avoids rounding fractional counts before import. Second, PRJNA747152 count matrices were pre-computed and loaded directly as integer counts with no minimum-expression prefilter and no length-aware import step.

#### 2.2.2 Proteomics

The two proteomics datasets (OSD-514, syn2580853) were preprocessed using distinct workflows according to their respective quantification strategies (TMT vs label-free). For OSD-514, Thermo RAW files from Orbitrap Fusion TMT runs were converted to mzML with ThermoRawFileParser and searched using FragPipe (v24.0, TMT10-MS3 workflow) against a target-decoy *D. melanogaster* UniProt database (UP000000803), with reporter-ion quantification referenced to a pooled control channel (ctrl_pool) across three TMT plexes (TMTA/B/C).

Peptide-spectrum matches were filtered at Q-value ≤ 0.01, PSM probability ≥ 0.90, and reporter-ion purity ≥ 0.50 (where available) and aggregated to protein-level abundances. Pool channels were excluded from downstream analysis. Proteins were included if quantified by ≥ 2 unique peptides, with positive finite abundances, and were present in all three TMT runs. Within each run, abundances were median-scaled, duplicate entries were summarized by median abundance, and only proteins with complete data across all samples were retained.

For syn2580853, label-free quantification (LFQ) intensities were extracted from MaxQuant protein output. Zero values were recorded as missing, and contaminant or reverse-database entries (CON, REV prefixes) were removed. Intensities were log2-transformed and proteins were retained if quantified in ≥ 70% of samples.

### 2.3 Differential Expression and Visualization

#### 2.3.1 RNA-seq

All three RNA-seq datasets were analyzed with DESeq2, using a negative binomial generalized linear model and Wald tests for pairwise contrasts. Raw p-values were adjusted for multiple testing with the Benjamini–Hochberg (BH) procedure, and differentially expressed genes were defined using an FDR-adjusted p-value (padj) below 0.05. A secondary effect-size threshold of |log2FC| ≥ 1 was applied when classifying genes as up- or down-regulated for volcano plots and gene-set selection. Log2 fold changes were shrunk with the apeglm estimator where a matching model coefficient was available; unshrunk estimates were retained for significance calls and downstream ranking.

Model designs and contrasts were dataset-specific. For OSD-514, the design formula was batch + sex + condition_group, reduced accordingly when batch or sex had only one level, with Earth as the reference level; three contrasts were tested (microgravity vs Earth, simulated 1g vs Earth, and microgravity vs simulated 1g), and ashr was used as a shrinkage fallback where a direct model coefficient was unavailable. For syn2580853, two models were fitted: an additive model (∼ day + genotype) testing the overall effect of Tau expression while adjusting for age, and a means model on a combined genotype–day factor (∼ group) from which timepoint-specific contrasts of TauWT vs Control at days 1, 10, and 20 were extracted. For PRJNA747152, the design was ∼ condition_group with Control (0 Gy) as the reference level, and five contrasts were tested (0.4 Gy vs Control, 10 Gy vs Control, combined 0.4 + 10 Gy vs Control, combined vs 10 Gy, and combined vs 0.4 Gy). Consistent with the absence of a minimum-expression prefilter for this dataset (Section 2.2.1), no batch term or sex covariate was included.

Results were visualized as volcano plots and heatmaps for each contrast. Volcano plots displayed log2 fold change against −log10 of the nominal p-value, with dashed guides at the padj < 0.05 and |log2FC| ≥ 1 thresholds; the most significant genes per contrast were labeled (ggrepel), and for PRJNA747152 the curated mitochondrial gene set (Section 2.4) was highlighted in place of generic top-gene labeling where mitochondrial genes were present. Heatmaps were generated from the top 50 genes by padj per contrast (40 for the timepoint-specific syn2580853 contrasts). Expression values were row-wise z-scored, clustered by gene, and displayed with samples held in a fixed condition order and annotated by experimental group; for syn2580853, z-scores were capped at ±2.5 to limit the influence of outlying samples on the color scale. For PRJNA747152, heatmaps were produced both genome-wide and restricted to the curated mitochondrial gene set. Plots were generated with ggplot2, ggrepel, and pheatmap.

#### 2.3.2 Proteomics

Differential protein abundance was assessed for both proteomic datasets in R with limma, fitting linear models to the processed log2 abundance matrices, computing moderated t-statistics by empirical Bayes shrinkage of the variance estimates, and adjusting p-values with the BH procedure.

For OSD-514, the primary model included condition, TMT run, and sex, and tested the contrasts simulated 1g vs Earth, microgravity vs Earth, and simulated 1g vs microgravity. Condition-only, condition-plus-batch, and sex-stratified models were additionally fitted as sensitivity analyses.

Significant proteins were defined by an adjusted p-value below 0.05 together with |log2FC| ≥ 1. For syn2580853, a means model was fitted on a combined genotype–day factor (∼ 0 + group), and nine contrasts were evaluated: TauWT vs Control, TauR406W vs Control, and TauR406W vs TauWT, each at days 1, 10, and 20. The mutant-versus-wild-type comparisons were treated as supplementary. Significant proteins were defined by an adjusted p-value below 0.05 together with |log2FC| ≥ 0.5, a lower effect-size threshold reflecting the more compressed dynamic range of label-free intensities relative to the TMT dataset.

Visualization followed the conventions described for the RNA-seq data. Volcano plots of log2 fold change against −log10 of the nominal p-value were generated per contrast with the most significant proteins labeled, and heatmaps were produced from the top 50 proteins per contrast, row-wise z-scored with samples ordered by condition and annotated by experimental group. For OSD-514, both uncorrected and batch-corrected PCA were generated from the processed abundance matrix.

### 2.4 Pathway and Functional Enrichment (GSEA)

For each pairwise contrast (SF1g vs Earth, SFμg vs Earth, SF1g vs SFμg), proteins were ranked using a signed significance score, computed as the sign of the log-fold-change multiplied by the -log10-transformed p-value, from the corresponding limma differential-abundance results, indexed by UniProt accession. GO Biological Process terms were retrieved for these UniProt IDs from org.Dm.eg.db (v3.21.0) via AnnotationDbi (v1.70.0), and gene sets were restricted to 10-500 members. Ranked lists were tested against these sets using fgseaMultilevel (fgsea v1.34.2), yielding normalized enrichment scores (NES) and BH adjusted p-values per term. Proteins meeting the differential-abundance significance thresholds (FDR < 0.05, |logFC| ≥ 1) were additionally mapped to gene symbols and submitted to Enrichr (FlyEnrichr) for GO Biological Process overrepresentation analysis, with top-enriched terms visualized as a term-protein concept network using igraph.

## 3. Results

### 3.1 Spaceflight-induced transcriptomic and proteomic changes (OSD-514)

Differential expression and pathway enrichment analyses of the OSD-514 dataset revealed distinct transcriptomic and proteomic responses under Earth, 1G spaceflight, and microgravity conditions (Figure 2). At the transcriptomic level, global volcano plots demonstrated a progressive shift toward transcript upregulation across the sequential pairwise comparisons from Earth to 1G spaceflight, and subsequently from 1G spaceflight to microgravity (Figure 2A) suggesting a cumulative transcriptional response to increasing departure from terrestrial gravitational conditions. However, pathway-level analysis revealed comparison-specific enrichment architecture rather than a uniform trend. Transcriptomic Gene Set Enrichment Analysis (GSEA) indicated substantial pathway enrichment in both the Earth-versus-1G-spaceflight and 1G-spaceflight-versus-microgravity comparisons, whereas the microgravity-versus-Earth comparison exhibited a more heterogeneous, mixed enrichment signature (Figure 2B). Of particular note, mitochondrial and energy-associated pathways including electron transport chain components and translation-related processes were preferentially enriched under microgravity relative to Earth conditions. Cytoplasmic translation was likewise enriched in microgravity relative to both 1G spaceflight and Earth, implicating a coordinated upregulation of protein synthetic machinery under conditions of gravitational unloading.

**Figure 2.**
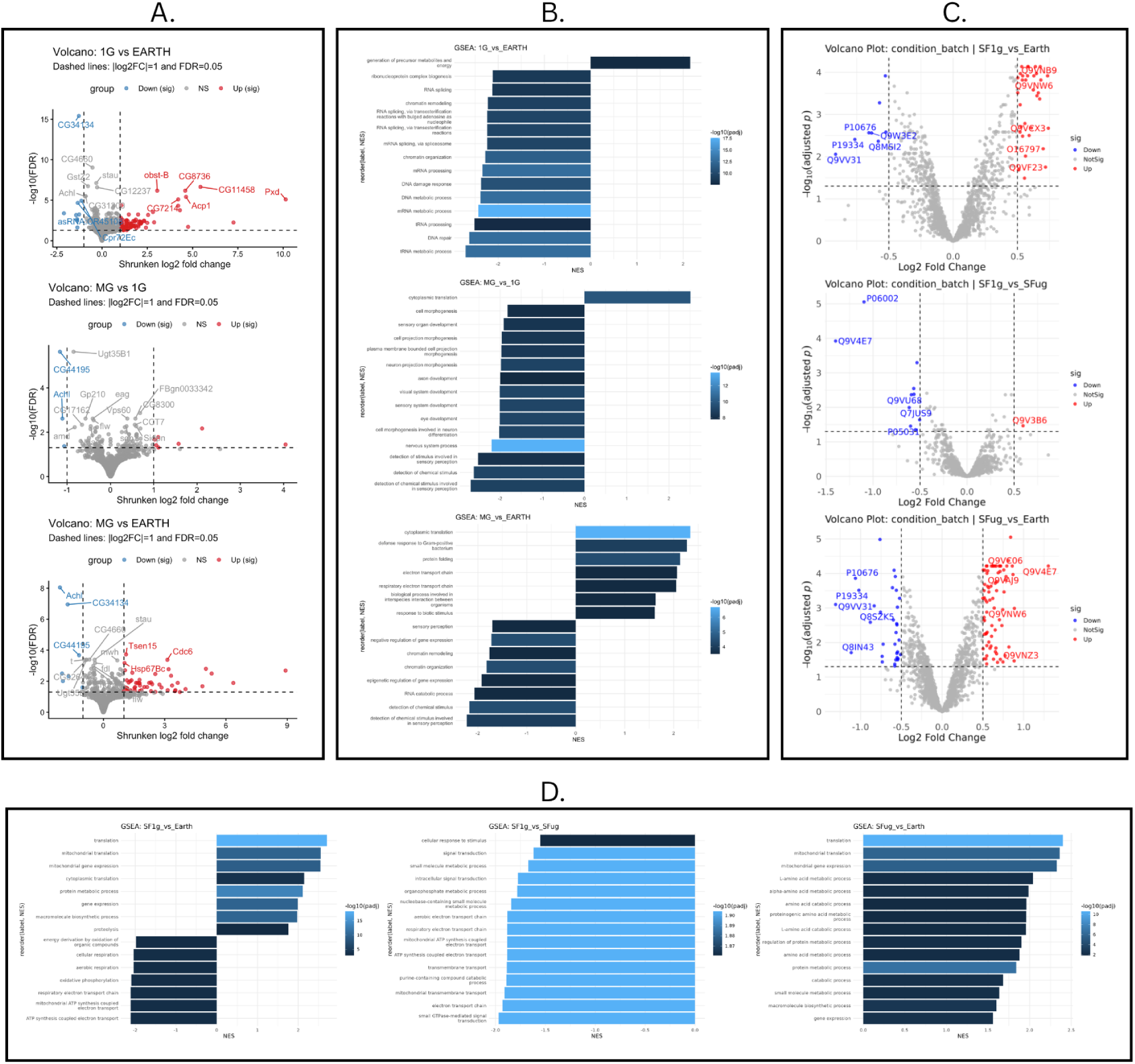
Transcriptomic and proteomic responses to spaceflight and microgravity in the OSD-514 dataset. (A) Global transcriptomic volcano plots comparing 1G spaceflight vs. Earth, microgravity vs. 1G spaceflight, and microgravity spaceflight vs. Earth, showing the predominance of upregulated transcripts across the sequential spaceflight conditions. (B) Transcriptomic Gene Set Enrichment Analysis (GSEA) comparing 1G spaceflight vs. Earth, microgravity vs. 1G spaceflight, and microgravity vs. Earth, highlighting comparison-specific enrichment of pathways related to mitochondrial function, energy metabolism, translation, and electron transport. (C) Global proteomic volcano plots comparing 1G spaceflight vs. Earth and microgravity vs. 1G spaceflight, showing a predominance of upregulated proteins with a more heterogeneous distribution than observed at the transcriptomic level. (D) Proteomic Gene Set Enrichment Analysis (GSEA) comparing 1G spaceflight vs. Earth, microgravity vs. 1G spaceflight, and microgravity vs. Earth, highlighting comparison-specific enrichment of pathways associated with mitochondrial respiration, ATP synthesis, electron transport, translation, gene expression, and protein metabolism.

At the proteomic level, volcano plot analysis similarly demonstrated a predominance of upregulated proteins across all pairwise comparisons, although this distribution was more heterogeneous than that observed transcriptomically (Figure 2C). Proteomic GSEA revealed more pronounced and consistent pathway enrichment specifically under microgravity conditions (Figure 2D). In the 1G-spaceflight-versus-Earth comparison, pathways governing aerobic electron transport, respiratory electron transport, and mitochondrial and non-mitochondrial ATP synthesis coupled to electron transport were relatively depleted under 1G spaceflight. At the same time, translation-associated pathways were concomitantly enriched in an apparent uncoupling of energy metabolism from protein synthetic capacity. By contrast, the microgravity-versus-1G-spaceflight comparison revealed enrichment of electron transport, ATP-related, membrane transport, and signal transduction pathways. Relative to Earth, microgravity was associated primarily with enrichment of translation, gene expression, catabolic, and broader metabolic pathways.

Collectively, these findings indicate that pathway-level enrichment was more pronounced under microgravity than under 1G spaceflight, particularly with respect to mitochondrial bioenergetics and energy-related processes. Across both the transcriptomic and proteomic datasets, recurrent enrichment of pathways governing electron transport, ATP production, translation, and cellular metabolism was observed, although the direction and magnitude of enrichment varied by comparison. These results suggest that the most robust, coordinated pathway-level perturbations within the OSD-514 dataset were elicited specifically under microgravity exposure, potentially reflecting an adaptive or compensatory bioenergetic response to the absence of gravitational loading.

### 3.2 Dose-dependent transcriptional response to gamma radiation (PRJNA747152)

GSEA (Figure 3A) of the PRJNA747152 dataset revealed a pronounced, dose-associated transcriptional response to gamma-radiation exposure. The Control (0 Gy) was the reference level and five contrasts, 0.4 Gy vs Control, 10 Gy vs Control, combined 0.4 + 10 Gy vs Control, combined vs 10 Gy, and combined vs 0.4 Gy, were tested. Transcriptional responses measured 8–10 h following exposure. The acute 10 Gy condition produced the most pronounced transcriptional response, indicating that radiation dose was associated with differences in the magnitude of the molecular response. Notably, the PCA graphs showed no clear separation by exposure condition among the Control, LowGamma, HighGamma, and ComboGamma groups, and volcano plot analysis identified relatively few significantly differentially expressed genes (Figure 3B). This suggests a comparatively modest gene-level transcriptional signature despite the dose-dependent pathway-level trends.

**Figure 3.**
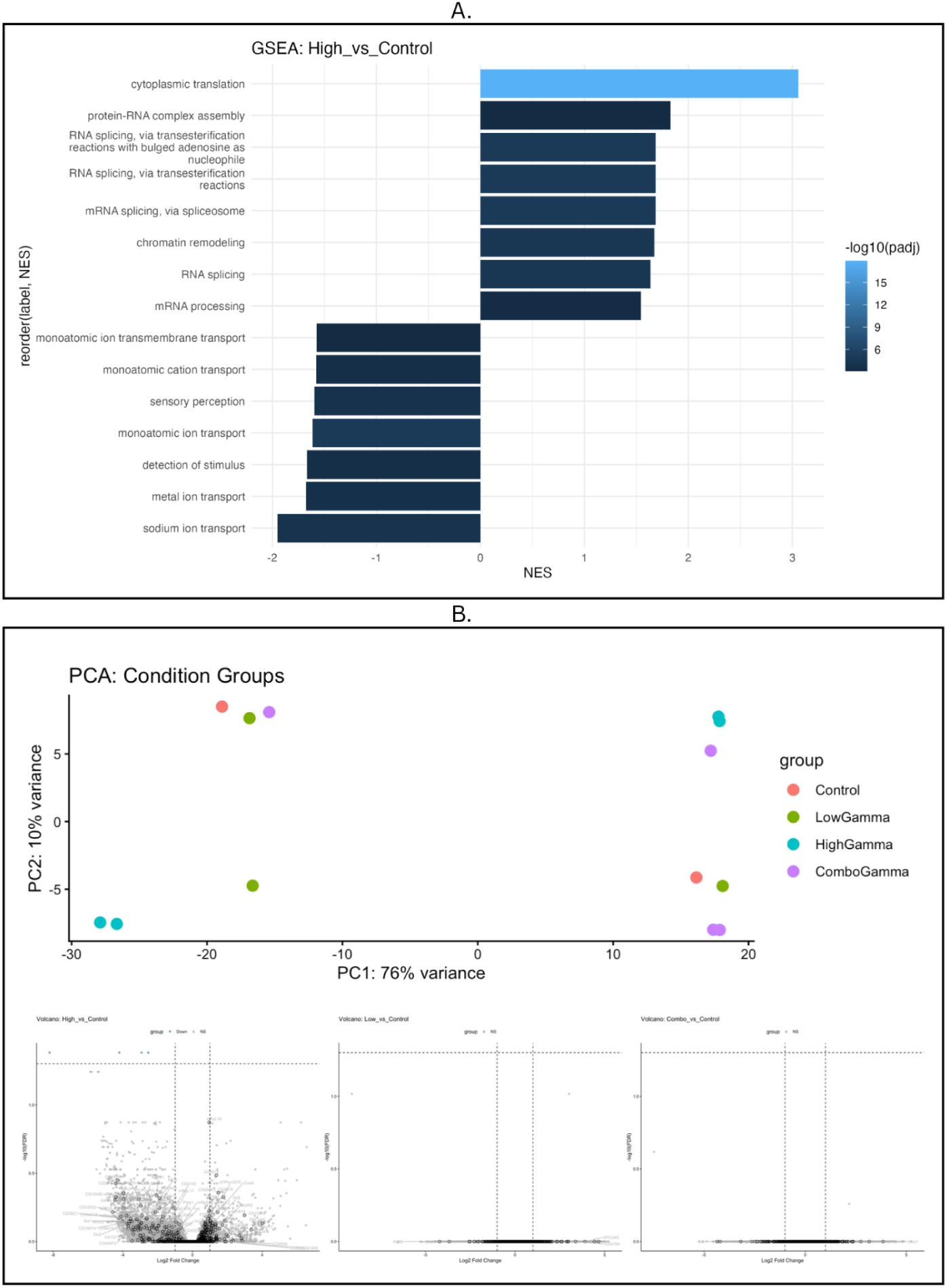
(A) Gene Set Enrichment Analysis (GSEA) comparing irradiated and control conditions, highlighting the biological pathways differentially enriched following gamma-radiation exposure. (B) Principal component analysis (PCA) of transcriptomic samples and global volcano plots comparing the Control, LowGamma, HighGamma, and ComboGamma groups. The PCA illustrates sample clustering according to exposure condition, while the volcano plots summarize differential gene expression across the radiation exposure groups.

The enriched pathways were predominantly associated with DNA damage response, oxidative stress, apoptosis, and broader cellular stress responses, consistent with the known biological effects of ionizing radiation. Higher-dose acute irradiation showed broader pathway enrichment than the lower-dose conditions, suggesting greater activation of stress-response programs with increasing radiation exposure. Oxidative-stress-related pathways were particularly relevant to the cross-condition analysis because they provide a plausible mechanistic point of convergence between radiation-induced cellular damage and the mitochondrial and redox-associated responses observed in the spaceflight and Tau datasets.

### 3.3 Tau-induced molecular remodeling (syn2580853)

Results show that Tau expression is associated with extensive transcriptomic and proteomic remodelling, with the neurodegenerative disease-associated R406W mutant producing substantially greater molecular disruption than wild-type Tau. R406W, WT and the control group were compared across days 1, 10 and 20. Proteomic principal component analysis (Figure 4A) demonstrated that TauR406W-expressing samples segregated more distinctly from controls than tau wild type, and there was pronounced separation evident between day one and day ten, while day ten and day twenty clustered more closely but remained distinguishable. This pattern is consistent with both temporal progression of the molecular phenotype and genotype-dependent differences that are sustained across the control, wild-type, and mutant conditions. Proteomic volcano plots (Figure 4B) further revealed a predominance in upregulated proteins at each time point. This occurred both in the comparisons between TauWT and the control groups on days 1, 10 and 20 and in the TauR406W vs Control Day20 groups. There was more differential gene expression in TauR406W vs Control than the WT vs Control.

**Figure 4.**
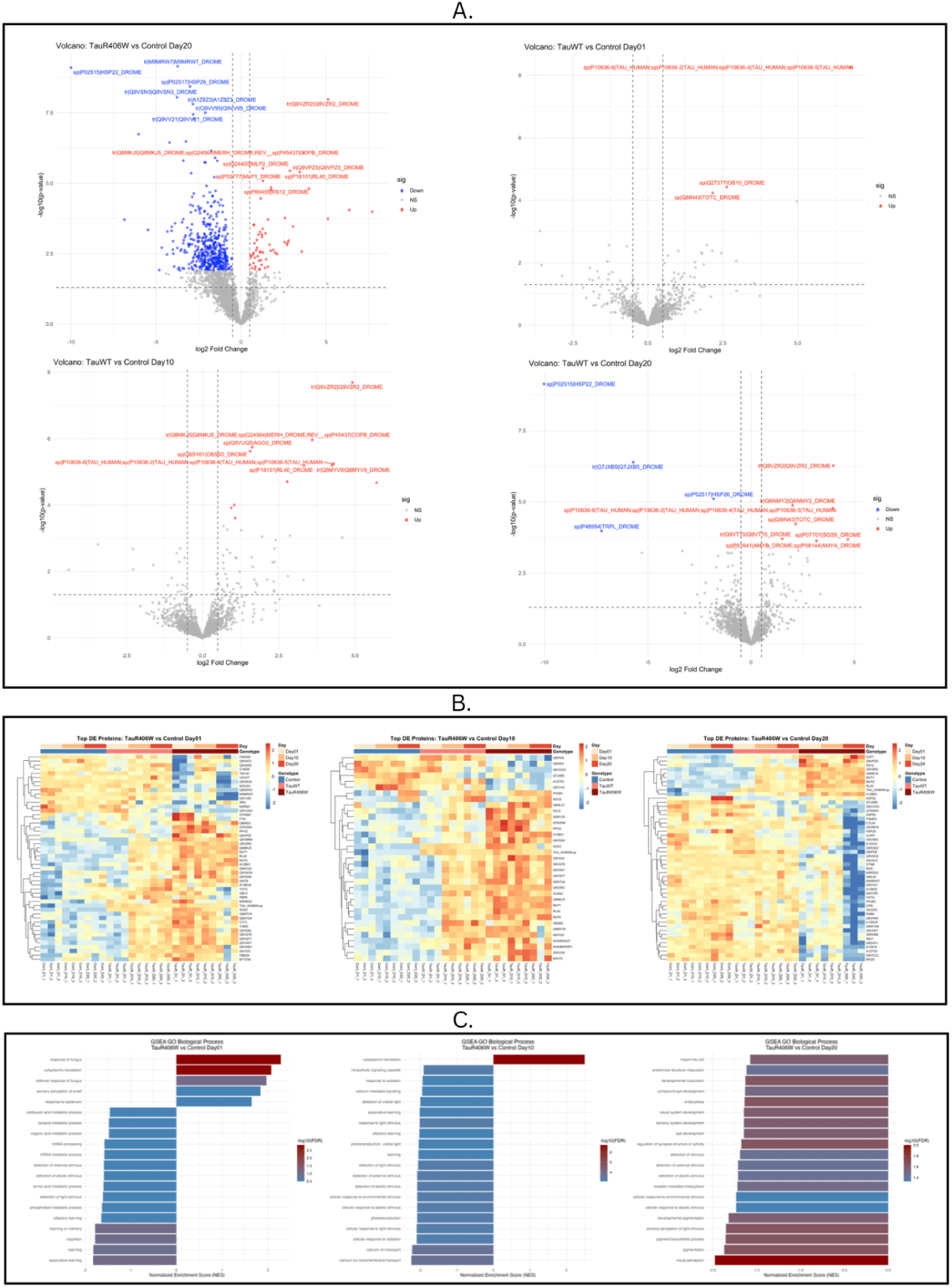
(A) Proteomic volcano plots for Days 1, 10, and 20 illustrating differential protein expression across the experimental groups. (B) Heatmaps of differentially expressed mitochondrial and oxidative stress-related proteins across Days 1, 10, and 20. (C) Gene Set Enrichment Analysis (GSEA) of biological processes for Days 1, 10, and 20 comparing genotype versus control, highlighting progressive alterations in mitochondrial and metabolic pathways, with Day 20 representing the strongest pathway-level changes.

Mitochondrial pathway analysis has shown the distinct effects of TauWT and TauR406W on energy related pathways. Heatmaps of the top differentially expressed proteins in TauR406W vs Control at Days 1, 10 and 20 (top), with GSEA GO Biological Process enrichment for each time point (bottom) are shown in Figure 4C. On day 20, relative to Control TauWT showed enriched electron transport, oxidative phosphorylation and ATP-synthesis pathways. This is distinct from from TauR406W where these mitochondrial energy-related pathways relative to tau wild type were depleted, which indicates a mutation-associated reduction in mitochondrial energy pathway enrichment. Membrane and vesicle-fusion pathways were also downregulated in both TauWT and TauR406W relative to control, with stronger depletion in TauR406W, suggesting that the mutation increases a deficit already present with the expression of Tau. The proteomic pathways for the aerobic electron transport chain, respiratory electron transport chain, mitochondrial ATP synthesis coupled electron transport, ATP synthesis coupled electron transport are all enriched in wild type relative to both control and R406W. However, this was enriched compared to Earth and not present in microgravity vs earth so it is unclear if this is a result of mutation. Gene Set Enrichment Analysis (GSEA) of biological processes (Figure 3D) for Days 1, 10, and 20 compared genotype versus control, highlighting progressive alterations in mitochondrial and metabolic pathways, with Day 20 representing the strongest pathway-level changes.

Taken together, these findings support mitochondrial dysfunction and synaptic impairment as central molecular consequences of pathological Tau accumulation, with the R406W mutation potentiating both phenotypes relative to wild-type Tau.

### 3.4 Cross-dataset convergence on mitochondrial signatures

To evaluate whether distinct environmental and genetic stressors converge on common mitochondrial molecular signatures, differentially expressed mitochondrial genes (DEGs) and proteins (DEPs) were compared across the OSD-514, syn2580853, and PRJNA747152 datasets (Figure 5). At the transcriptomic level, overlap between datasets was minimal, with only a single mitochondrial DEG (CG10814) shared between the spaceflight and Tau datasets and no genes shared across all three datasets. The PRJNA747152 dataset showed no overlap with either OSD-514 or syn2580853, indicating that the transcriptional response to gamma radiation was largely dataset-specific and did not converge with spaceflight- or Tau-associated transcriptional signatures at the individual gene level.

**Figure 5.**
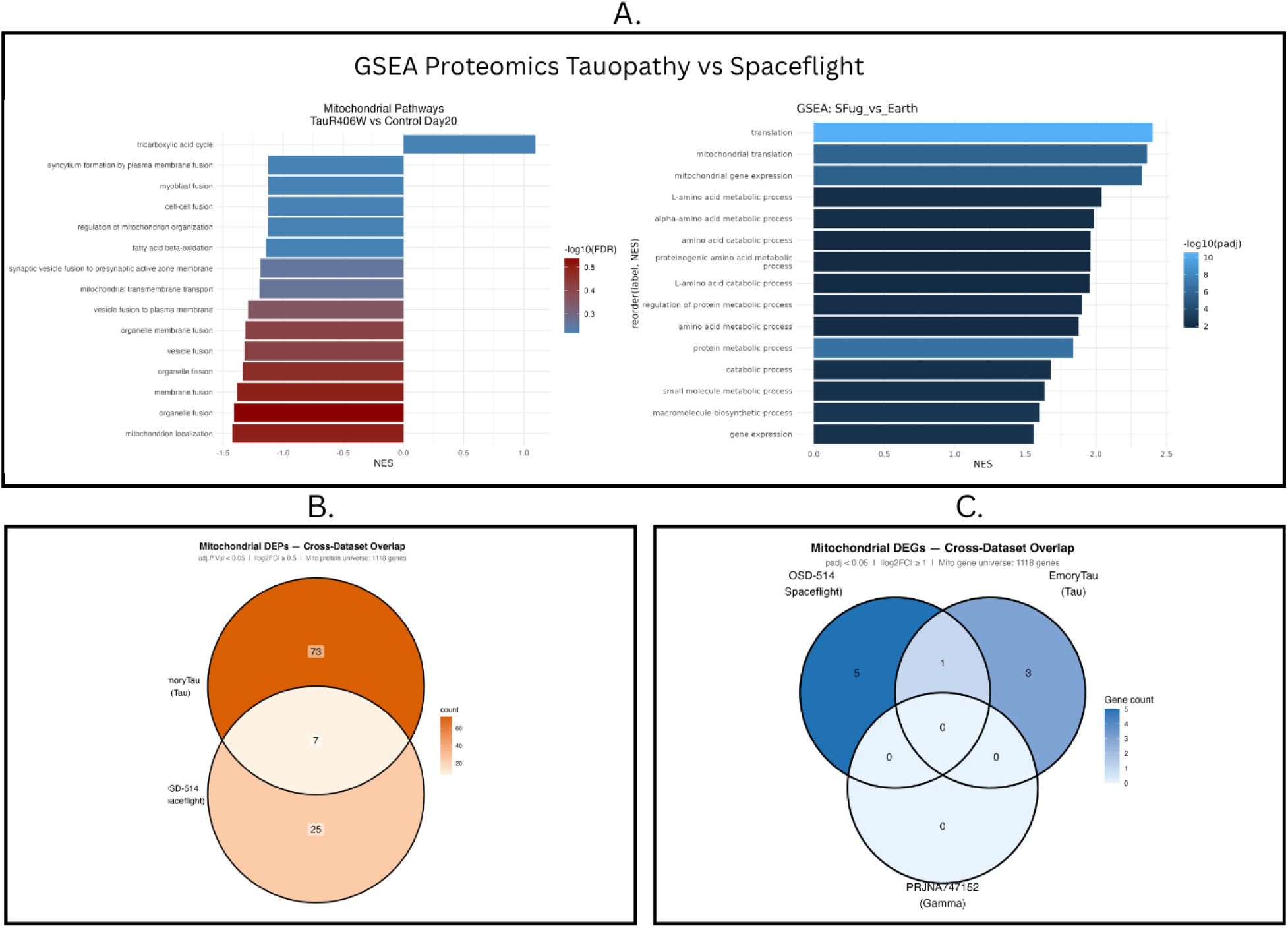
(A) Comparative Gene Set Enrichment Analysis (GSEA) of the representative proteomic comparisons from the OSD-514 (SFμg vs. Earth) and syn2580853 (Tau R406W vs. Control, Day 20) datasets. Shared mitochondrial pathways, including electron transport chain and ATP synthesis-related processes, were identified in both datasets. Because enrichment was calculated using different reference groups, pathway directionality cannot be directly compared between datasets. (B) Venn diagram showing the overlap of mitochondrial differentially expressed genes (DEGs) across the OSD-514 (spaceflight), syn2580853 (Tau), and PRJNA747152 (gamma radiation) datasets. Only a single mitochondrial DEG (CG10814) was shared between the spaceflight and Tau datasets, whereas no genes overlapped with the gamma-radiation dataset or across all three datasets. (C) Venn diagram showing the overlap of mitochondrial differentially expressed proteins (DEPs) between the OSD-514 and syn2580853 datasets. Seven mitochondrial DEPs were shared between both datasets, while PRJNA747152 was not included because proteomic data were unavailable.

Proteomic comparison, by contrast, revealed substantially greater convergence than was observed transcriptomically. Seven mitochondrial differentially expressed proteins (DEPs) were shared between the OSD-514 and syn2580853 datasets; proteomic comparison with PRJNA747152 was not feasible, as proteomic data were unavailable for this dataset. The shared proteins aldh, drp1, Mpcp2, mtpbeta, pdgy, sdhA, and sfxn1-3-exhibited opposing directionality between the two datasets, being upregulated under spaceflight conditions but downregulated in the Tau model. This inverse relationship may reflect divergent adaptive versus pathological mitochondrial responses to distinct stressors converging on a shared set of protein targets.

## 4. Discussion

This study compared whole-brain transcriptomic and proteomic responses in *D. melanogaster* across microgravity, Tauopathy and ionizing radiation to test whether the spaceflight-associated stressors and AD-associated stressors converge on shared mitochondrial signatures. Microgravity was associated with coordinated upregulation of mitochondrial electron transport and energy-related pathways. Interestingly, tau expression showed the opposite pattern, with a suppression of oxidative phosphorylation (OXPHOS) and ATP synthesis pathways. Gamma radiation exposure produced a stress response with minimal gene-level change. Taken together, these findings show that convergence between spaceflight and Tauopathy was minimal at the transcript level but more substantial at the protein level, and these shared proteins changed in opposite directions between conditions. We had hypothesized that spaceflight and Tauopathy would produce the same mitochondrial response, making spaceflight a faster model of AD-related dysfunction. Instead, the two conditions responded in opposite directions, suggesting that spaceflight-stressed organisms mount a compensatory response that tau-expressing organisms cannot. Identifying where compensation gives way to failure may reveal early markers of neurodegeneration and inform crew health monitoring on long missions.

### 4.1 Differentially Expressed Genes Across Datasets

The differential expression analysis identified only one significant gene, CG10814, in the comparison between the microgravity and Alzheimer’s disease (AD) Tauopathy conditions. Despite the limited number of significant transcript-level changes, CG10814 exhibited the same overall directional pattern observed in the broader analysis, being upregulated under microgravity and downregulated in the AD Tauopathy condition. This opposing regulation suggests that, although microgravity and Tauopathy produced limited overlap at the individual gene level, they may induce contrasting responses in mitochondrial-associated processes.

The identification of only a single significant gene is particularly notable when considered alongside the proteomic results. The limited transcriptional response contrasts with the broader differences observed at the protein level, indicating that changes in transcript abundance alone may not fully capture the molecular response to microgravity. In particular, the discrepancy between the RNA-seq and proteomic analyses suggests that processes occurring downstream of transcription may contribute to the observed changes in protein abundance. These processes could include translational regulation, protein degradation, and mechanisms affecting protein stability or activity. However, because these mechanisms were not directly assessed in the present analysis, this interpretation remains hypothetical.

The presence of a stronger signal at the proteomic level also demonstrates the importance of integrating multiple molecular datasets when investigating mitochondrial responses to spaceflight. If the analysis had been restricted to differential gene expression, the relatively small number of significant transcripts could have suggested a limited molecular response. The proteomic analysis instead revealed broader alterations, indicating that transcriptional changes may not necessarily correspond directly to changes at the protein level.

### 4.2 Proteomics Convergence and Functional Breakdown of Core Mitochondrial Hubs

Unlike the results of the transcriptomic analysis which identified only a single shared gene between spaceflight and Tau datasets, proteomics analysis revealed a significant convergence between those datasets, identifying the shared mitochondrial differentially expressed proteins (DEPs) aldh, drp1, Mpcp2, mtpbeta, pdgy, sdhA, and sfxn1-3. Importantly, all seven of these proteins demonstrated a bimodal expression pattern, displaying significant upregulation under isolated microgravity conditions but faced downregulation in the Tau pathology model. GSEA further confirmed these observations through broader group effects that mirrored the DEP patterns, tying the influence of microgravity to core mitochondrial processes including OXPHOS and ATP synthesis. Examining the specific roles of each of these seven shared proteins reveals key aspects of mitochondrial bioenergetics regulation relevant to the understanding of Alzheimer’s pathology:

Pyruvate dehydrogenase (pdgy) regulates the conversion of cytosolic pyruvate into acetyl-CoA, functioning as the primary metabolic gatekeeper that connects glycolysis to the Citric Acid Cycle (TCA) [51-53]. Upregulation during microgravity reflects an increase in TCA activity whereas its suppression in tauopathy restricts the entry of acetyl-CoA, reducing the production of ATP and creating an energy deficit [53-55].

Succinate dehydrogenase complex subunit A (sdhA) makes up the catalytic subunit of Complex II in the ETC and supports the flow of the TCA cycle by converting succinate into fumarate [56]. Downregulation of this protein in tauopathy not only disrupts the ETC, but also negatively impacts the TCA, both of which results in reduced ATP generation [56-58]. Its upregulation we observed in microgravity may indicate a cellular compensatory response to maintain mitochondrial function.

Regulation of Phosphate Import for ATP coupling (Mpcp2/SLC25A3 ortholog): Mitochondrial phosphate carrier protein (Mpcp2) is responsible for importing inorganic phosphate across the inner mitochondrial membrane into the mitochondrial matrix [59-60]. There, Complex V (ATP synthase) of the ETC combines it with ADP to form ATP [61]. Our finding that Mpcp2 is decreased in tauopathy suggests impaired mitochondrial phosphate import, which would be expected to reduce ATP synthase activity and contribute to the broader OXPHOS deficit in tau pathology. In tauopathy, ETC disruptions such as increased mitochondrial reverse transport are further associated with increased leakage of electrons and a build-up of ROS, further amplifying bioenergetic stress [62].

SFXN1 and SFXN3 are likely mitochondrial transmembrane serine importers while SFXN2 regulates iron and heme homeostasis [63-65]. Because serine metabolism supports redox balance and heme serves as a prosthetic group for ETC complexes II, III, IV, and Cytochrome C, dysregulation of these SFXN proteins likely results in impaired mitochondrial metabolism and neuronal vulnerability consistent with neurodegeneration [64, 67-68].

Dynamin-related protein 1 (drp1) regulates mitochondrial fission and helps control the distribution of mitochondria in neurons [69-71]. Downregulation of drp1 in syn2580853’s tauopathy model is consistent with prior studies showing that hyperphosphorylated tau disrupts DRP1-dependent mitochondrial dynamics through abnormal interactions, resulting in impaired mitochondrial transport, increased oxidative stress, and reduced neuronal bioenergetic capacity [72-74].

Aldehyde dehydrogenase (aldh) detoxifies reactive aldehydes derived from ROS-driven lipid peroxidation [75-76]. Current research links impaired ALDH activity in tauopathy and Alzheimer’s disease to accelerating mitochondrial dysfunction, ROS accumulation, tau hyperphosphorylation in vulnerable neurons, and even sensitizing neurons to glutamate-induced cytotoxicity [43, 77-78].

Mitochondrial trifunctional protein subunit beta (mtpbeta) catalyzes the final step of long-chain fatty acid degradation to produce Acetyl-CoA for use in the TCA [35]. Impaired oxidation of fatty acids associated with tauopathy leads to lipid accumulation within astrocytes and microglia, fueling the release of pro-inflammatory cytokines that contributes to neuroinflammation and secondary oxidative stress [18, 35, 79].

Across all seven protein hubs, we observed a statistically significant upregulation from microgravity in contrast to a pronounced downregulation in the tauopathy model. Additionally, GSEA enrichment results mirror this bi-directional protein expression, providing robust multi-level validation that these proteins serve important roles in the overall governance of OXPHOS and ATP synthesis. This multi-omics level convergence reinforces that microgravity and tauopathy act on identical bioenergetic pathways but through opposing effects: microgravity stressor triggers a compensatory response that boosts bioenergetics pathways that are otherwise impaired and downregulated in tauopathy.

While transcriptomic analysis yielded only a single shared differentially expressed gene, proteomic analysis revealed significant convergence between microgravity (OSD-514) and the Tauopathy model (syn2580853), identifying seven shared core mitochondrial proteins: aldh, drp1, Mpcp2, mtpbeta, pdgy, sdhA, and sfxn1-3. Notably, each shared protein demonstrated an inverted expression profile where there was a decrease in protein levels within Tauopathy but a reciprocal increase under microgravity. GSEA confirmed these findings at the pathway level, strengthening the findings that microgravity and tau pathology act on identical bioenergetic machinery but through opposing mechanisms.

#### 4.2.1 Substrate Gating and Bioenergetic Coupling

We observed that pdgy/PDH, sdhA/SDHA, Mpcp2/SLC25A3, and mtpbeta/HADHB were decreased in the Tauopathy model and increased in microgravity. Given the established roles of PDH in linking glycolysis to acetyl-CoA production [52-53], HADHB in mitochondrial fatty acid beta-oxidation [80], SDHA in coupling the TCA cycle to the electron transport chain through Complex II [56, 81], and Mpcp2 in importing inorganic phosphate for ATP synthesis [59], this pattern we observed is consistent with a shift in mitochondrial substrate utilization contributing to impaired OXPHOS in Tauopathy. Specifically, reduced abundance of these proteins in the Tauopathy model likely impairs bioenergetic capacity, whereas the reciprocal increase in abundance observed within microgravity implies a compensatory response aimed at maintaining bioenergetic pathways. However, the compensatory nature of this response remains inferential unless further supported by functional or direct flux assays.

#### 4.2.2 Quality Control, Antioxidant Defense, and Dynamic Remodeling

A similar reciprocal regulation was also observed among aldh/ALDH, sfxn1-3/SFXN1–3, and drp1/DRP1. Based on existing literature, ALDH is thought to detoxify reactive aldehydes generated during lipid peroxidation [76] while SFXN proteins maintain mitochondrial competency through serine transportation in the ETC [63, 82]. The reduction of these protein levels within the Tauopathy model thus suggests a reduction in oxidative-stress defense in neurons [64]. For drp1/DRP1 specifically, while our Tauopathy model observed decreased protein abundance, the broader literature indicates that tau pathology drives dysregulated mitochondrial fission through abnormal Drp1 activity [83] rather than just a simple reduction in Drp1 abundance alone: Drp1 interacts abnormally with hyperphosphorylated tau [72] and alters Drp1 localization to mitochondria [74]. However, the decrease in Drp1 abundance we observed might still result in functional abnormalities and ultimately contribute to mitochondrial fission dysregulation. Similar to the bioenergetic pathways discussed before, inferences about altered functional mitophagy, fission dynamics, or ROS clearance using only protein abundance data remains limited, and requires additional validation using live-cell imaging, ROS quantification, and organelle turnover assays.

### 4.3 Radiation Specificity Control

In the weightlessness vs 1G comparison, the influence of microgravity was isolated and observed to yield a significant difference in DEG and DEP between the two groups. However, in addition to microgravity, cosmic radiation remains a chronic primary stressor for cells in space. To analyze the isolated effects of radiation on bioenergetic pathways, we introduced the PRJNA747152 dataset, which revealed a dose-dependent stress-response pattern associated with activation of mechanisms tied to DNA damage, apoptosis, and oxidative-stress. Although ionizing radiation did not directly modulate bioenergetic pathways, both microgravity and radiation converge on elevated oxidative stress, a key driver of neuroglial network damage and a well-established hallmark of neurodegeneration.

Inclusion of the PRJNA747152 dataset helped isolate the effects of acute ionizing radiation on cellular responses, revealing a selective increase in repair- and stress-response pathways without affecting mitochondrial bioenergetics. However, it should be noted that the absence of a bioenergetic effect could be attributed to species-specific radioresistance in Drosophila that may not translate directly to human biology [84-85]. Additionally, even though the PRJNA747152 dataset evaluated acute ionizing radiation, which is distinct from the chronic radiation exposure in space, the combination of both OSD-514’s internal controls for chronic radiation along with PRJNA747152’s pathway findings further supports microgravity as being the primary driver of the observed bioenergetic boost.

Finally, while microgravity, cosmic radiation, and Tauopathy are all known to induce elevated oxidative stress, PRJNA747152 showed that radiation increases oxidative stress through distinct pathways separate from microgravity or Tauopathy. DNA damage from radiation can occur through direct physical ionization from high-energy photons, however up to 70% of the damage could be indirectly attributed to radiolysis [86-87]. Instead of striking the DNA molecule directly, high-energy photons convert neighboring water molecules into ROS that diffuse and chemically alter nearby DNA molecules, resulting in acute oxidative stress. This likely resulted in the radiation dose-dependent effects we observed, where radiation-induced oxidative stress drove DNA damage repair mechanisms and neuronal-microglial apoptosis. On the other hand, microgravity-induced oxidative stress is likely tied to mitochondrial remodeling and altered ROS regulation while Tauopathy-related oxidative stress results from damaged bioenergetics pathways leaking ROS.

### 4.4 Glial-Neuronal Metabolic Decoupling

Neurons have a high metabolic baseline yet maintain relatively low intrinsic antioxidant defense, requiring astrocytic support for metabolic needs and microglia function for maintaining homeostasis. As such, neuronal mitochondrial pathophysiology associated with neurodegeneration is considered non-cell-autonomous. Under normal physiological conditions, astrocytes maintain neuronal bioenergetics through several key mechanisms: The Astrocyte-Neuron Lactate Shuttle (ANLS), Astrocytic Fatty Acid Degradation, and Intercellular Mitochondrial Transfer. In the ANLS, astrocyte transport proteins export the glycolytic energy substrate lactate into neurons, which are then converted into pyruvate within the TCA [93-95]. To prevent lipotoxicity in neurons, astrocytic mitochondria undergoes fatty acid oxidation to remove peroxidized lipids [35, 96]. During bioenergetic stress, glia transfer functional mitochondria to neurons to support OXPHOS and maintain energy needs [88].

Our research identified 7 key protein hubs that were upregulated under microgravity, of which 3 are directly tied to these key astrocytic processes: mtpbeta/HADHB, sfxn1-3/SFXN1-3, and drp1/DRP1. HADHB helps catalyze beta-oxidation of fatty acids in the mitochondria, but in neurodegeneration where fatty-acid breakdown is impaired [80], lipids start to build up in astrocytes and contribute to secondary oxidative stress [97]. Sideroflexins, especially SFXN1 with supporting function from SFXN3, maintain mitochondrial competency by importing serine across the inner membrane for mitochondrial one-carbon metabolism, with likely downstream effects on glycine-dependent heme synthesis [63, 82]. Although SFXN2’s role is less clear, it is thought to carry out similar functions as SFXN1 and SFXN3 [63]. Critically, SFXN1 plays an important role in maintaining complex III functionality in the ETC [82]. In healthy glia, maintaining a pool of fully competent mitochondria is essential for non-cell-autonomous organelle transfer to support neuronal bioenergetics [38, 40]. In our dataset, sfxn1-3/SFXN1-3 was one of the seven core mitochondrial hubs that exhibited reciprocal regulation. The observed reduction in SFXN1-3 protein abundance in the tauopathy model suggests a decrease in glial mitochondrial competency, implying a depleted pool of functional mitochondria available for export to neurons. Meanwhile, Drp1-mediated fission is required prior to packaging mitochondria for intercellular transport [43]. Although mammalian AD models often show elevated Drp1 activity [73], *Drosophila* tauopathy models exhibit mitochondrial Drp1 mislocalization [74], which could explain the decrease in protein abundance we observed.

Additionally, previous research has linked excessive Drp1-mediated mitochondrial fission to neuronal degeneration in mice models [43, 89], further implicating the impact of Drp1 dysregulation. For both Drp1 and sideroflexins, the core translational insight is that tau pathology likely leads to model-dependent Drp1 dysregulation converging on mitochondrial dysfunction, impairing glial-neuronal bioenergetics coupling and contributing to oxidative stress and neurodegeneration.

SFXN1-3 proteins are key for maintaining mitochondrial competency. Under normal conditions, glia export healthy mitochondria to neurons to support their bioenergetics, however when under inflammation or neurodegenerative stress, glia have been observed to transfer damaged mitochondria to neurons instead [38, 40, 90]. In the context of our study, we identified a significant decrease in SFXN1-3 expression within the Tauopathy model, suggesting a potential decrease in functional mitochondria that could be exported to neurons and highlights the possibility of damaged mitochondria being absorbed as well, both of which decreases energy production in neurons while the latter may increase oxidative stress.

Drp1 coordinates mitochondrial fission to prepare mitochondria for vesicular transport to neurons. As with SFXN1-3 and HADHB, Drp1 was also observed to be significantly downregulated in the Tauopathy model while being significantly upregulated under microgravity. However, existing research remains conflicted on whether Drp1 remains increased or decreased in AD, as it appears to be model and context-dependent. For fruit flies, Tauopathy appears to deplete mitochondrial Drp1 through mislocalization, which matches our study’s observations [74]. With diminished levels of Drp1, less mitochondria will be able to be shuttled to neurons, once again limiting energy production in neurons. Overall, while the literature supports the idea that Drp1 disruption is conserved across Tauopathy models, the direction of Drp1 impact appears context-dependent, with mammalian Tauopathy models displaying increased Drp1 expression and mitochondrial fission activity [73]. Therefore the strongest translational point from using a Drosophila model is the conserved dysregulatory impact on mitochondrial dynamics in the context of AD.

### 4.5 Cross-Species Generalization and Translational Relevance

Although *D. melanogaster* remains an efficient model for multi-omics testing under various conditions, generalizing findings to mammalian biology requires accounting for species-specific differences. It should be noted that while physiological differences do exist, several fundamental bioenergetic pathways such as the ETC [91], TCA cycle [92], and orthologous proteins such as sfxn1-3 [63] are remarkably conserved across species. However, specific regulatory processes may still diverge, as in the case with Drp1 whose protein abundance and activity levels in *Drosophila* Tauopathy models do not necessarily reflect those seen in mammals. Still, rather than quantifying identical protein changes across species, the value behind the *Drosophila* model lies in its ability to identify conserved OXPHOS control points, such as bioenergetics substrate gating through pdgy/PDH and mtpbeta/HADHB or mitochondrial competency through sfxn1-3/SFXN1-3. Additional testing of the inverted proteomic results using rodent spaceflight models and potentially human iPSC-derived neuronal-glial cells can further validate whether the microgravity-induced bioenergetic compensation mechanisms we observed in *Drosophila* translates to mammalian systems.

### 4.6 Limitations

These findings should be taken with several limitations in mind. First, sex differences could not be assessed in this analysis. Despite containing both male and female samples, the Tauopathy dataset (syn2580853) was pooled and did not distinguish between the two. Furthermore, the ionizing radiation dataset (PRJNA747152) only contained male samples. Although sex was modeled as a covariate in OSD-514, the absence of comparable data across all three datasets precludes any cross-condition inference about sex.

Second, proteomic data was unavailable for the radiation dataset, so protein-level convergence could be assessed only between spaceflight and Tauopathy. Given the greater overlap observed at the protein level than at the transcript level, our conclusions about radiation may underestimate its mitochondrial effects. This is compounded by the radiation tolerance of D. melanogaster. Young adult flies have reported LD50 values of about 1,238 Gy for males and 1,339 Gy for females [84], and another gamma-irradiation study reported an LD50 of about 840 Gy [85]. Because the doses in PRJNA747152 (up to 10 Gy) fall far below these thresholds, the minimal gene-level response may reflect species-specific tolerance rather than an absence of radiation effects on mitochondria. The radiation results should therefore be interpreted as specific to this organism and dose range. Given that the radiation dataset contributed little to the cross-condition overlap, we expect this limitation had a modest effect on the core spaceflight-Tauopathy comparison.

Finally, the exposures examined here are graded, but each was sampled at only a few levels. Specifically, these were three gravity conditions in OSD-514 (Earth 1g, simulated 1g, microgravity), and two radiation doses in PRJNA747152 (0.4 and 10 Gy). Dose- or gravity-dependent trends in the mitochondrial response therefore could not be observed. The radiation doses used here are well above those present aboard the ISS, therefore the radiation dataset cannot be treated as a direct, dose-matched proxy for the radiation component of spaceflight.

### 4.7 Future Directions

Future work should include sex-stratified analyses in datasets with identifiable male and female samples, and proteomic profiling of ionizing radiation-exposed tissue. Functional assays of respiration, mitochondrial membrane potential, ROS, and mitophagy in microgravity and Tauopathy models would test the compensation hypothesis. Combined exposures that reflect the spaceflight environment would clarify whether early mitochondrial upregulation gives way to dysfunction. Finally, validation in mammalian models and human tissue would verify if these findings are unique to *D. melanogaster*, as well as establish translational relevance.

## Acknowledgements

The authors thank the NASA Biological and Physical Sciences (BPS) Division for supporting the NASA Open Science Data Repository (OSDR), including NASA GeneLab and the NASA Ames Life Sciences Data Archive (ALSDA). We thank the OSDR team for curating the OSD-514 (MVP-Fly-01) dataset and GeneLab for processing the RNA-seq data, and we thank Dr. Sharmila Bhattacharya and colleagues for generating the original MVP-Fly-01 transcriptomic and proteomic data (Mhatre et al., 2022). This work was conducted through the OSDR Brain Analysis Working Group and its Mitochondria Subgroup Project.

The results published here are in whole or in part based on data obtained from the Accelerating Medicines Partnership for Alzheimer’s Disease (AMP-AD) Target Discovery Consortium data portal and can be accessed at doi:10.7303/syn2580853", as well as the Sequence Read Archive (NCBI) under BioProject PRJNA747152.

## Author Contributions

**Marian Navarro**: Conceptualization, Methodology, Software, Formal Analysis, Writing – original draft, Writing – review and editing, Visualization, Project administration **Jelard Aquino**: Software, Formal Analysis, Writing – original draft **King Shi**: Software, Formal Analysis, Writing – original draft **Henry Lin**: Writing – original draft **Vahid Monfared**: Writing – original draft **Joven Christopher Garcia**: Writing – original draft **Allen Seylani** Writing – original draft **Maria Komędera** Writing – original draft **Julia Moutinho** Writing – original draft **Jessica Romils** Writing – original draft **Andrea Venosa** Writing – original draft **Anish Kelam** Writing – original draft **Vandreza Bortolotto** Writing – review and editing **Jingchun Chen** Funding acquisition, Resources, Supervision, Conceptualization, and Writing – review & editing **Matthew Gebre** Writing – review and editing **M Windy McNerney**: Validation, Writing – review and editing, Supervision, Project Administration

## Notes

### Competing Interest Statement

The authors have declared no competing interest.

