## Supplemental Figure 1 for "Convergent Mitochondrial and Oxidative Stress Signatures in Spaceflight and Tauopathy: A Comparative Multi-Omics Reanalysis in Drosophila melanogaster"

### OSD-514 Transcriptomics PCA

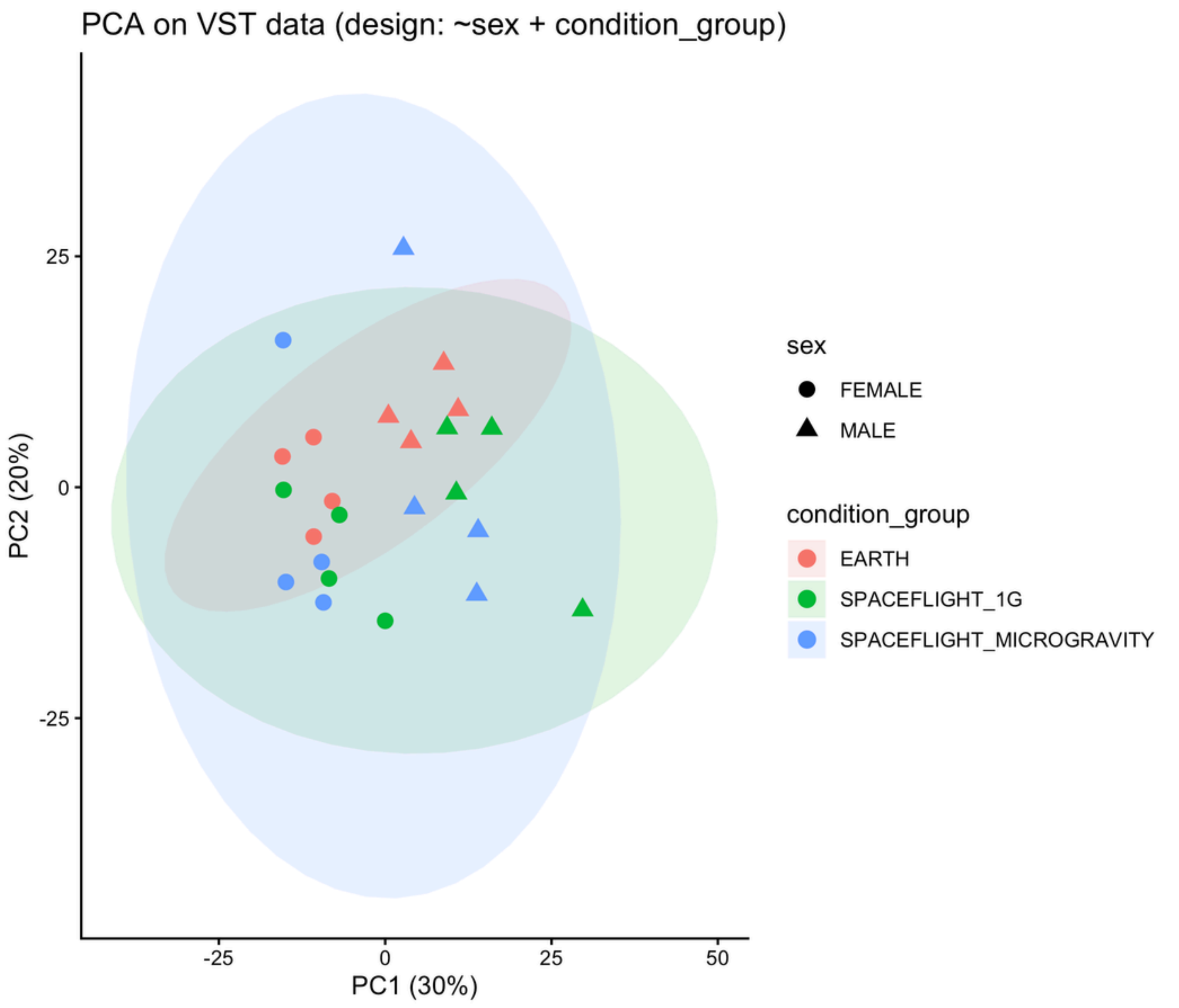


### OSD-514 Proteomics PCA

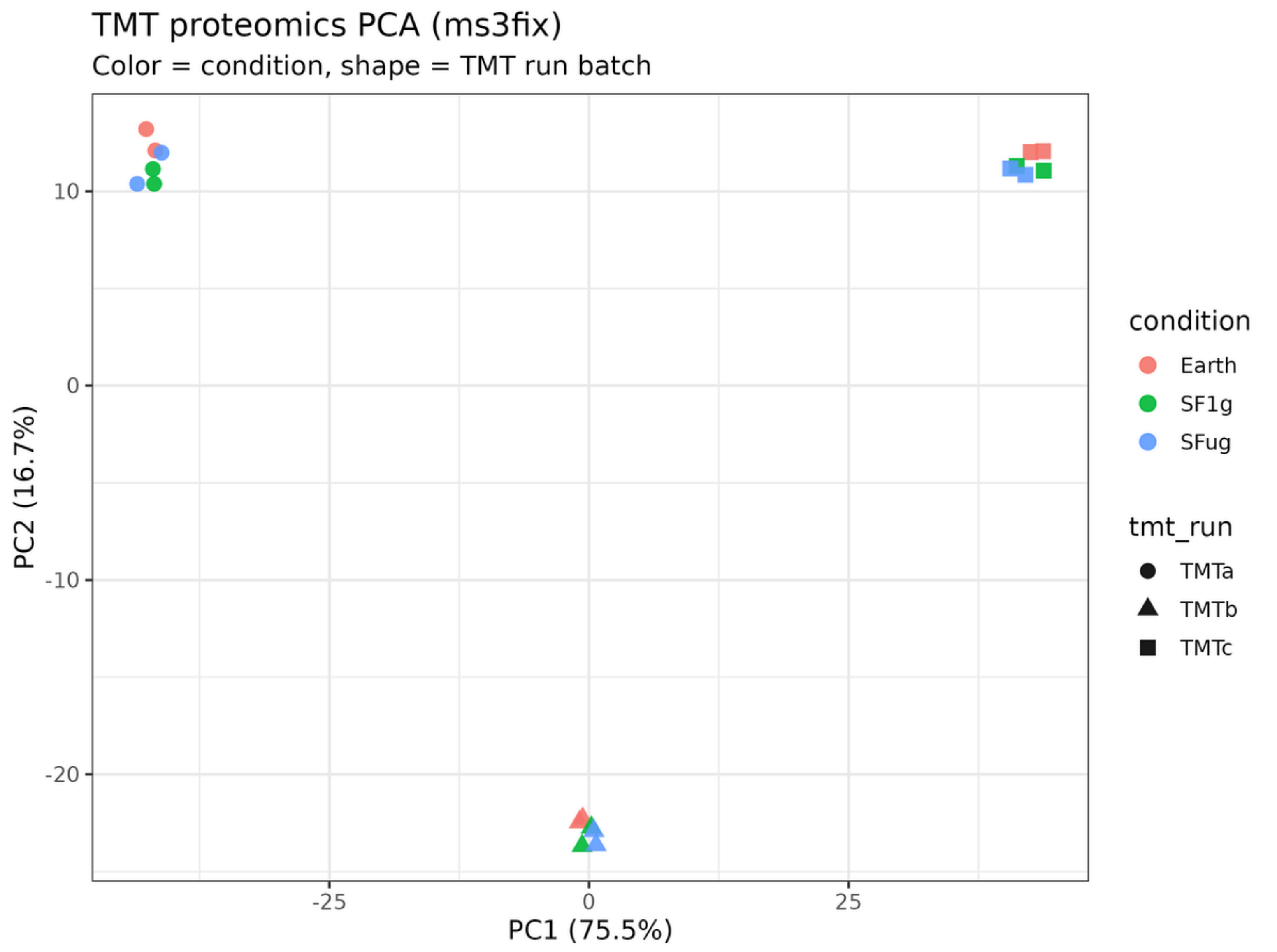


### syn2580853 Transcriptomics PCA

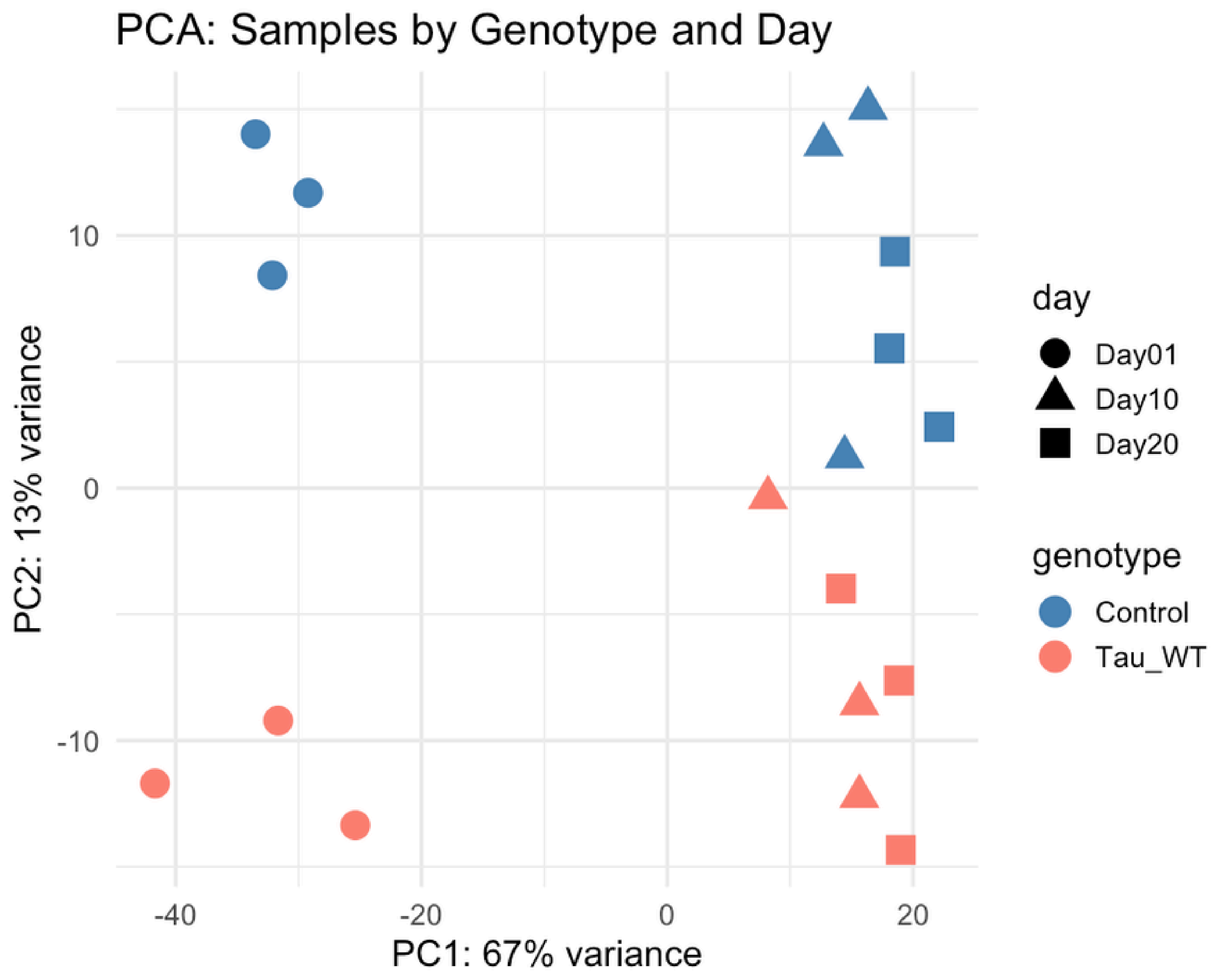


### syn2580853 Proteomics PCA

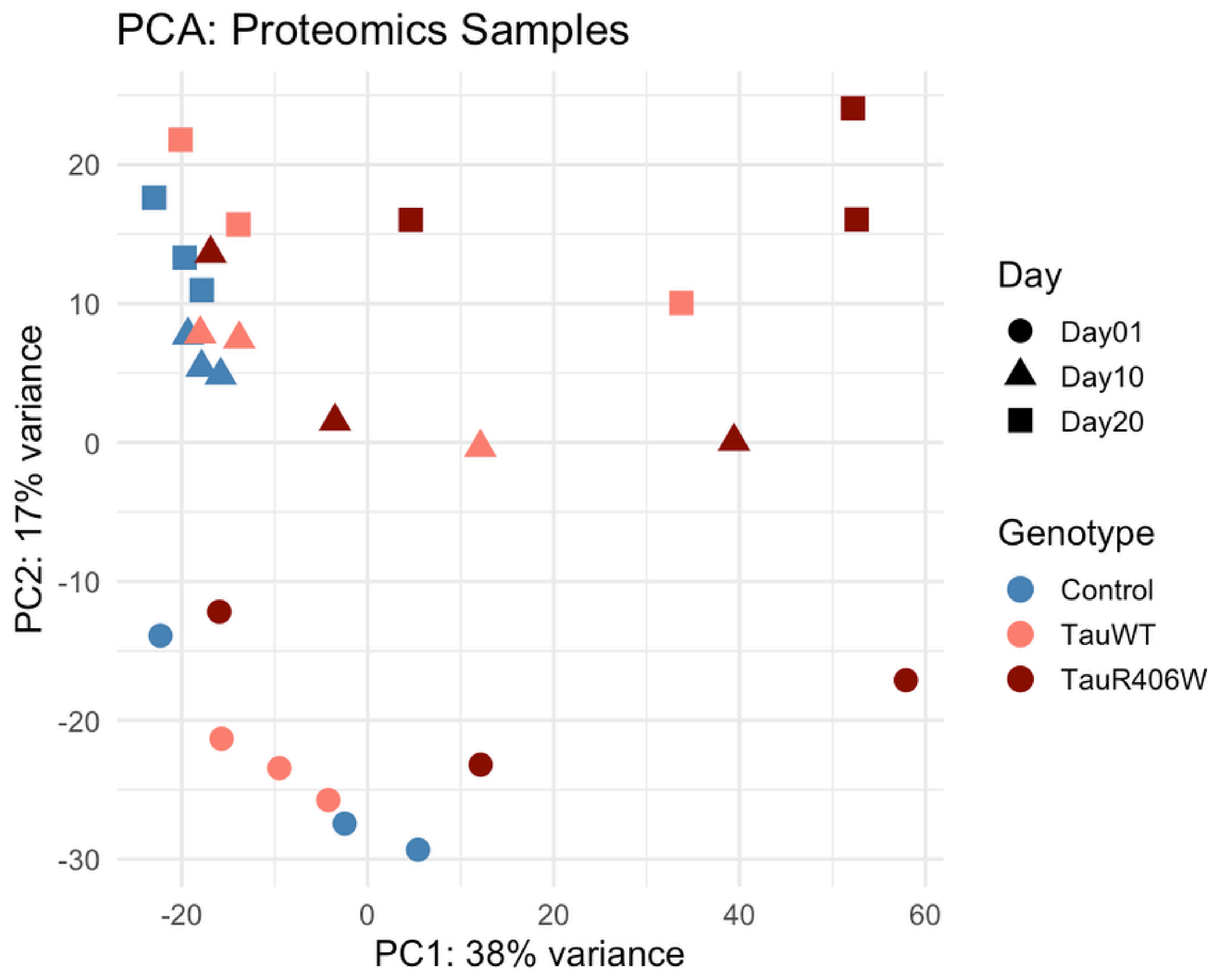


### PRJNA747152 Transcriptomics PCA

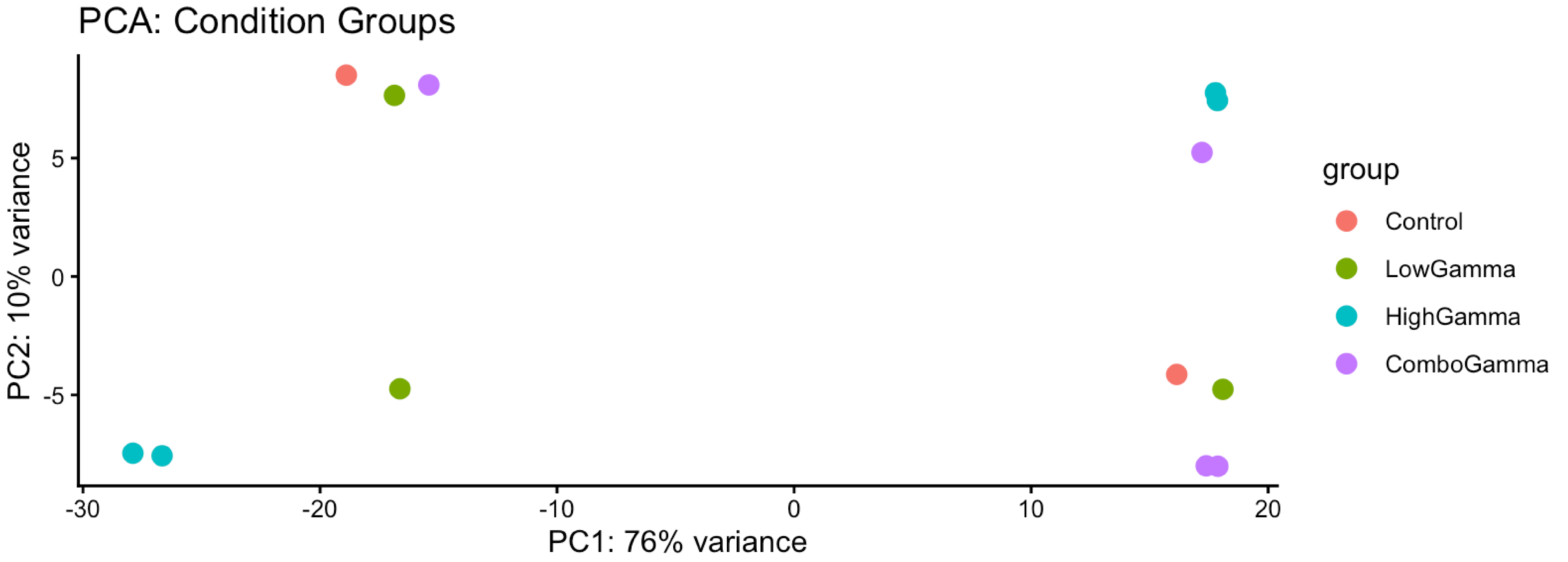
